# Grasp Aperture Corrections in Reach-to-Grasp Movements Do Not Alter Size Perception

**DOI:** 10.1101/2021.02.18.431812

**Authors:** Vonne van Polanen

## Abstract

When grasping an object, the opening between the fingertips (grip aperture) scales with the size of the object. If an object changes in size, the grip aperture has to be corrected. In this study, it was investigated whether such corrections would influence the perceived size of objects. The grasping plan was manipulated with a preview of the object, after which participants initiated their reaching movement without vision. In a minority of the grasps, the object changed in size after the preview and participants had to adjust their grasping movement. Visual feedback was manipulated in two experiments. In experiment 1, vision was restored during reach and both visual and haptic information was available to correct the grasp and lift the object. In experiment 2, no visual information was provided during the movement and grasps could only be corrected using haptic information. Participants made reach-to-grasp movements towards two objects and compared these in size. Results showed that participants adjusted their grasp to a change in object size from preview to grasped object in both experiments. However, a change in object size did not bias the perception of object size or alter discrimination performance. In experiment 2, a small perceptual bias was found when objects changed from large to small. However, this bias was much smaller than the difference that could be discriminated and could not be considered meaningful. Therefore, it can be concluded that the planning and execution of reach-to-grasp movements do not affect the perception of object size.

## 1 INTRODUCTION

Sensory information can be used to guide our movements and to perceive the world around us. For example, when grasping an object, visual information about object size can be used to scale the opening between the fingertips (grip aperture) to the size of the object [1]. Grasping an object will also give haptic information about object size which can be optimally integrated with vision to obtain an accurate percept of object size [2]. In an influential theory, the dual-stream theory, it has been suggested that processing of visual information for guiding actions and perception are separated in two streams in the brain [3].

Since the introduction of the dual-stream theory of action and perception, there has been discussion about the separation between the two streams. Support for this theory has been provided with behavioural experiments, for example in the grasping domain, where it has been proposed that the perception of object size and the planning and execution of grasping is dissociated. This was specifically shown with visual illusions, where objects appeared to perceptually differ in size based on the visual context, while grasp apertures were not affected by the illusion [4, 5].

However, there has been much controversy about the nature of this dissociation between grasping and size perception (see for reviews [6, 7]). In contrast to the initial studies, later studies showed effects of illusionary size on both size perception and grasp parameters [8, 9]. Several explanations have been proposed to explain these contradictory findings. For instance, it has been argued that an illusion effect on grasping depends on the availability of visual feedback during grasping [10] or that the effect is only apparent in planning but not execution [11]. Furthermore, an alternative view from a separation between processing of action and perception is a separation between absolute versus relative measures [12]. These authors argue that in the previous studies, perception of object size was a relative comparison of two sizes, whereas for grasping an absolute measure of size is used which is not affected by an illusionary context. However, if task context is manipulated, there appears to be no difference between action and perception per se, but between whether relative or absolute measures are needed.

While currently it is acknowledged that perception and action processes are not completely separate, but interact with each other (see e.g. [13–15] for reviews), there are still few studies that specifically address the interactions between these processes. Especially the literature on effects of actions on perception is scarce. Some studies found that perception was improved depending on the action context. Gutteling, Kenemans (16) showed that orientation change detection was improved depending on the subsequent action plan. Similarly, reaction times in a visual search task were altered depending on the subsequent action that had to be executed [17].

Furthermore, a few other studies showed that perception can also be altered by the action context. For instance, there are studies that have shown effects of grasping context on size perception or graspability judgements. Graspability refers here to the ability to grasp an object of a specific size with one hand: if the object is too large, it cannot be grasped with one hand. When potential grasping ability was altered, e.g. by increasing the apparent hand size, size perception was altered as well ([18], but see also [19]). Similarly, the judged ability whether a participant could grasp an object was altered when changing grip aperture by squeezing a ball or spreading the fingers [20]. While this study found an effect on graspability judgements, it did not find effects on size judgements. It is possible that a complete reach-to-grasp movement compared to a static alteration of grip aperture could be more effective in altering size perception. Such an effect was shown by Cesanek and Domini (21), who found that visuo-haptic adaptation of the grasping movement in a virtual reality setup did influence manual estimation of object size.

All in all, it seems that action context can alter the perception of objects. However, it is not clear how planned actions and potential corrections to these action plans could be incorporated into the perceptual judgement. Interestingly, in a study on object lifting, it was found that planned fingertip forces altered the perceived object weight [22]. When an object was lighter than expected, forces had to be decreased to skilfully lift the object. It was found that objects were also perceived to be lighter and this change in weight perception correlated with the decrease in force scaling. A similar effect was found when lifting objects with an asymmetric weight distribution. The torque planning error was associated with an increase in weight perception and torque estimation [23]. This suggests that a correction to an action plan could induce a change in perception in a similar direction as the correction.

In object lifting, fingertip forces have to be controlled and scaled towards the object weight. Corrections to the applied force will often rely on haptic feedback of perceived weight. In reach-to-grasp movements, different motor parameters have to be controlled (i.e. finger positioning) and visual feedback can also be used to implement corrections. Therefore, it is not clear whether corrective movements could also influence perception in reach-to-grasp movements. The present study used a reach-to grasp task to investigate whether corrective actions to grasping could influence size perception. Specifically, two questions were addressed: 1) whether corrections to action plans would also affect perception in reach-to-grasp tasks similar to object lifting tasks, and 2) what would be the role of visual and haptic feedback in these corrections.

To address these issues, a task was used in which participants grasped different objects and compared these in size. They were presented with a preview of an object in order to plan a grasping movement. Next, they initiated the reaching movement without vision. In some trials, the object would be changed, so they had to correct their movement. Several studies have shown that grasp apertures can be adjusted online when an object is suddenly changed after the movement has been initiated [24–27]. It was investigated whether these movement corrections would affect the perception of object size. The availability of visual and haptic feedback was manipulated in two experiments. In Experiment 1, visual information was provided after the initiation of the movement and could be used to correct the grip aperture. In Experiment 2, vision was not available and only haptic feedback of the object could be used to correct the grip aperture. In accordance with van Polanen and Davare (22), who found that downscaled force corrections were associated with a reduction in perceived weight, a similar directional effect is hypothesized for the present study: if the grip aperture is decreased because the object is smaller than expected, it is hypothesized that objects will be perceived to be smaller.

## 2 METHODS

### 2.1 Participants

Thirty right-handed participants took part in the experiment. They all had normal or corrected-to-normal vision and no known neurological impairments. Fifteen participants performed Experiment 1 (age 24±4 years, 11 females) and the other fifteen took part in Experiment 2 (age 24±4 years, 10 females). They all signed informed consent before participating. The experiments were approved by the local ethical committee of KU Leuven.

### 2.2 Apparatus

Movements were measured with a TrakSTAR system (NDI). Two sensors were placed on the nails of the thumb and index finger. Sensor positions and orientations were captured with a sample frequency of 240 Hz. Participants wore liquid crystal goggles (PLATO, Translucent Technologies) goggles to control their vision. The glasses could be in an opaque or transparent state, which was controlled by a NI-USB 6343X (National Instruments) connected to a personal computer.

The objects to be grasped were plastic black bars of different sizes. The width and height of the bar was 2 cm, while the length varied. A small and a large set was used with a reference length of 4 and 8 cm, respectively. The small set ranged from 2.75-5.25 cm with steps of 0.25 cm. A smaller step size of 0.125 was used between 3.5 and 4.5 cm, giving a total of 13 bars in the small set. This smaller step size enabled a more precise measure of size differences around the reference value. Similarly, the large set consisted of 13 bars ranging from 6.75-9.25 cm. The stimuli are shown in Figure 1A.

**Figure 1.**
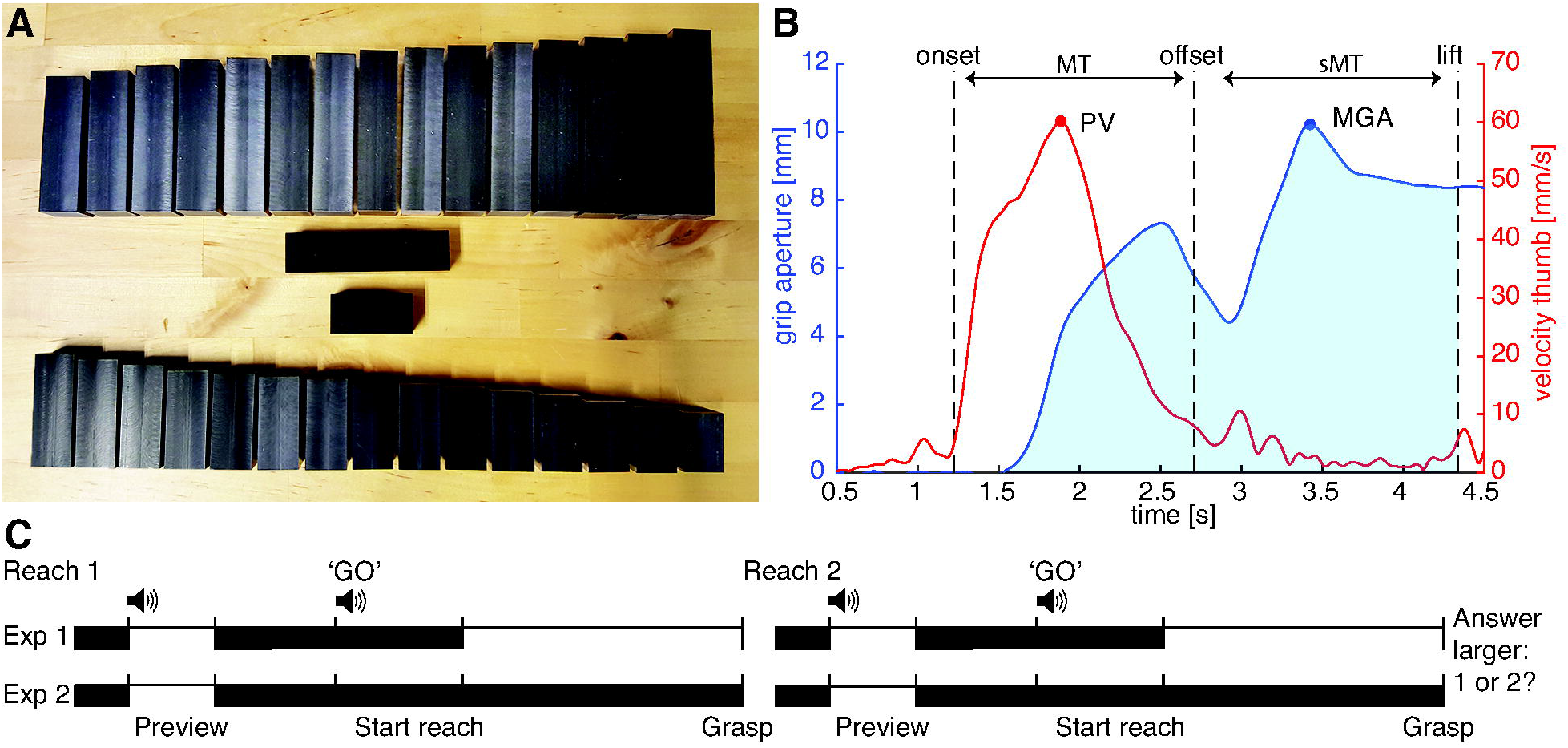
**A**: Stimuli in the large (top) and small set (bottom), with the two reference sizes in the middle. **B**: Example of a single reach-to-grasp movement in Experiment 2, in the correct large condition. Note that this is an example with a large correction in grip aperture for illustrative purposes; many trials also had more smooth corrections. Solid lines indicate grip aperture (blue) and thumb velocity (red). Dashed lines represent time points of movement onset, movement offset and start of the lift, respectively. The parameters of interest are indicated: maximum grip aperture (MGA), peak velocity (PV), movement time (MT), secondary movement time (sMT) and summed grip aperture (shaded area). Note that the summed grip aperture was calculated over the curve plotted against the relative time (see Figure 4). **C**: Experimental time-line in Experiment 1 and 2 of 1 trial, consisting of 2 reaches. White and black parts indicate availability of visual information, i.e. when goggles are open and closed, respectively. First, a preview is shown with a warning tone. Next, after participants hear the ‘GO’ signal, they can initiate their reach-to-grasp movement. Vision is restored only in Experiment 1. After two reaches, participants indicate which of the two grasped objects was larger.

### 2.3 Procedure

Participants sat in front of a table and wore the goggles. They performed reach-to-grasp movements where they moved from a starting position to one of the stimulus bars, grasped it, lifted it briefly and placed it back on the table. The starting position was marked with a round soft padding (±1 cm width), so participants could locate it easily without looking. The bar was positioned ±42-49 cm from the starting position, adjusted to a comfortable reaching distance for each participant. The reach-to-grasp task consisted of two phases: a preparation and an execution phase. In the preparation phase, participants kept their hand at the starting position with index finger and thumb pinched together. A beep sounded to alert them and the goggles opened briefly (500 ms) and showed them a stimulus bar placed at the target position. They were told to use the information in the preparation phase to plan their movement. After this preview, the goggles closed again and participants waited for the start signal to perform the reach-to-grasp movement in the execution phase. In the execution phase, they should reach for the bar and grasp it. The start of the movement was indicated with a verbal ‘go’-signal provided by the computer. When they initiated their reaching movement, vision was still blocked by the goggles. In Experiment 1, visual feedback was given after movement initiation until the end of the execution phase. More specifically, the goggles opened after a distance of ± 6 cm was covered by the marker on the thumb (on average 290 ms after movement onset) and stayed open for 5 seconds. In Experiment 2, no visual information was given during the execution phase and the goggles stayed closed throughout the execution phase. This second experiment was performed to examine the role of visual and haptic feedback and make sure a grasping movement was needed to evaluate the size of the object. That is, in the first experiment, participants could have judged object size based on visual information without the need to grasp the object. In the second experiment, the grasping movement was necessary to judge the size (i.e. contacting the object to obtain haptic information) which could increase a potential effect of the grasping movement on the perceived size. An outline of both experiments is given in Figure 1 C. Participants were instructed to move at a comfortable speed, but grasp the bar in one fluent motion.

Sometimes the bar changed in size between the preparation and the execution phase. In this case, the experimenter replaced the bar after the preview. A soft cloth was placed on the table, so the changing of objects was not audible to the participants. In addition, even if the object was not changed, the original object was lifted and replaced by the experimenter at the same position to make sure any auditory cues were present in each trial. A change in object size was performed only in 25% of the reaching movements (50% of trials, which always had 2 reaching movements), so participants were expected to still rely on the visual information in the preparation phase. The bars were always placed at the same location, so participants knew where the object was and did not have to search for it.

Participants always performed two sequential reach-to-grasp movements in one trial (one reference and one test object). After the second grasp, they were asked to indicate which of the two grasped bars was larger. They were instructed to compare the bars they grasped, not the ones during the preview. The reference objects were the 4 cm and 8 cm bars. The test objects were bars from the large or small set. Note that only reaches for reference objects could contain a change and test objects never changed in size. Therefore, the preview of a test object was always the same as the grasped test object. When the reference object was not changed (no correction needed), the preview and grasped object were of the same size, i.e. 4 or 8 cm. When the reference object changed in size (correction needed), the other reference size was shown in the preview: for example, when the object changed from large to small, the 8 cm bar was shown in the preview and the 4 cm bar had to be grasped. Therefore, the grasps of reference objects were varied in 4 conditions: no-correct small (4-4), correct small (8-4), no-correct large (8-8) and correct large (4-8). Here, the first number refers to the previewed object and the second number to the grasped object. In the correct small and no-correct small conditions, the grasped reference object was the small, 4 cm, object and the test objects from the small set were used. For example, a trial could consist of an 8-4 reference and a 5.25-5.25 test grasp. In the correct large and no-correct large conditions, the large, 8 cm, object was used as grasped reference and the test objects from the large set were used. For example, a trial could have a 4-8 reference and a 9.25-9.25 test grasp. In this way, the reference object that was grasped was compared to test objects similar in size. In the no-correct conditions, objects did not change between the preparation and execution phase. In the correct large and correct small conditions, the reference object changed from small to large and from large to small after the preview, respectively. The large size difference between the preview and the grasped object in the correct conditions (4 cm vs. 8 cm) was chosen to have clear differences in planned grasping movements and induce large corrective movements.

The order of trial presentation was determined by using a staircase procedure. For each condition, two staircases were used: one starting with the smallest and one starting with the largest object in the test set. For example, the no-correct small condition started with a test object of 2.75 cm in one staircase, and a test of 5.25 cm in the other. When participants judged the test object to be larger than the reference object, the test object was decreased in size for the next trial. When the test object was judged to be smaller, it was increased in size for the next trial. The 8 staircases (two for each condition) were presented in an interleaved manner. Staircases were terminated after 7 reversals. Since it was possible that participants expected the object to change if they saw a large object in the second preview after grasping a small object (or vice versa), a 9^th^ series was interleaved among the staircases, where 4 cm and 8 cm object grasps were performed, with a preview of the same size as the grasped object. This 9^th^ series was terminated after 10 trials. For all trials, the order of test and reference grasps was randomized. On average, 136 and 131 trials were performed in Experiment 1 and 2, respectively. A short break was provided after approximately 50 trials. An example of the staircases from a representative participant is shown in Figure 2.

**Figure 2.**
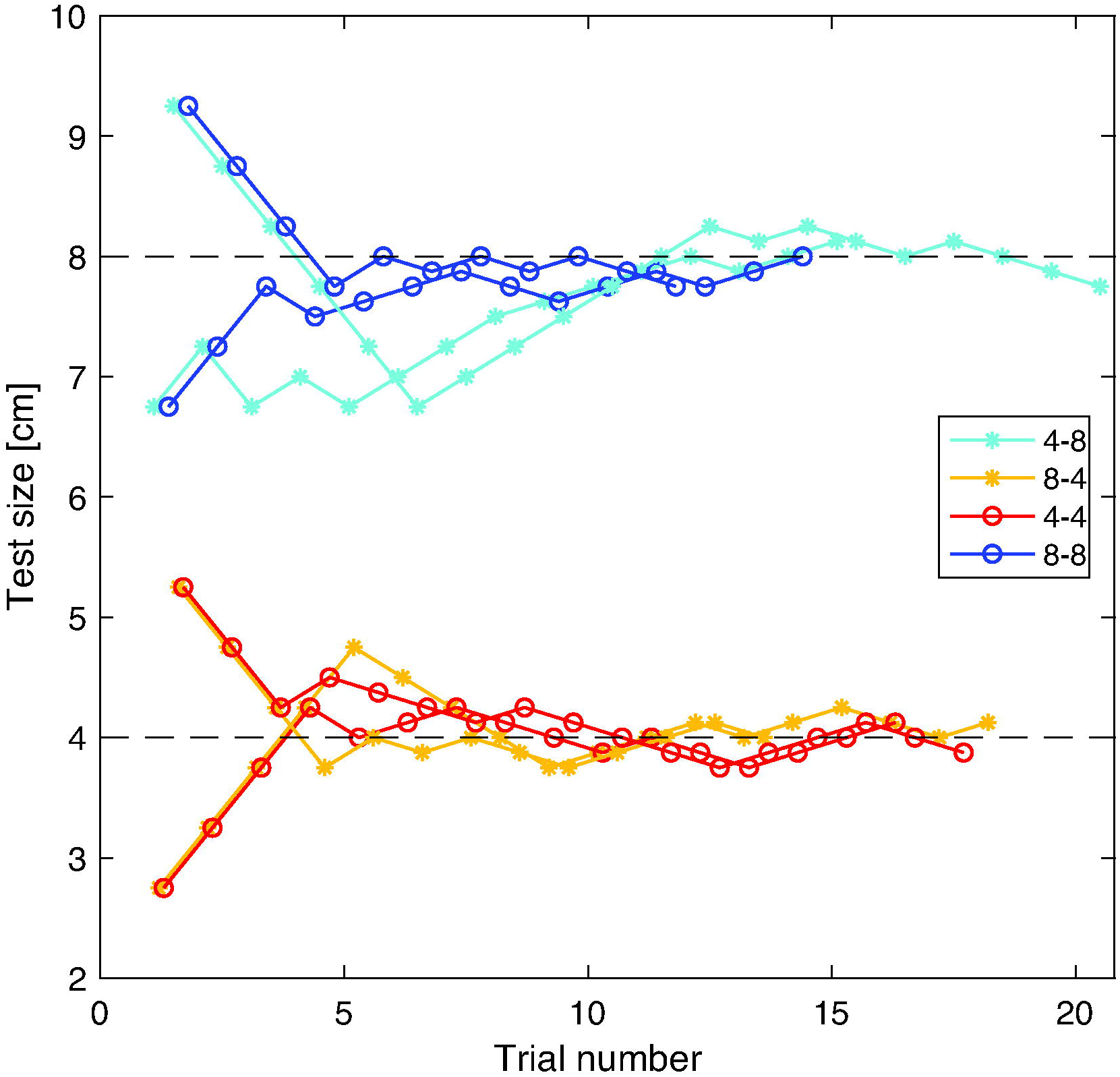
Example of staircase procedure for a representative participant in Experiment 1. Each condition has two staircases, one starting at a high and one at a low value in the test set. Circles represent no-correct conditions (4-4: no-correct small and 8-8: no-correct large) and asterisks indicate correct conditions (8-4: correct small and 4-8: correct large). The 9^th^ series with 4 and 8 cm objects is not shown here. Dashed lines indicate the size of the reference objects.

### 2.4 Data analysis

#### 2.4.1 Perceptual answers

Answers from the staircases in the four conditions were plotted against the test size as the percentage of trials in which the test was judged to be larger than the reference. Psychometric curves were fitted through these percentages, using the following equation:

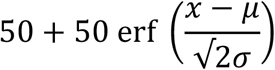

where *μ* represents the point of subjective equality and *σ* the standard deviation, which is a measure of the discrimination threshold. A weighted fit was used to account for the different number of trials for different test sizes. Biases were calculated by subtracting the reference value from the point of subjective equality and were expressed as a percentage difference from the reference size. Similarly, the *σ* was divided by the reference size and expressed as a percentage to obtain the weber fraction. This is the relative difference that can be discriminated. By taking the relative values, values for the small and large set could be compared more easily.

#### 2.4.2 Movement analysis

To compare reach-to-grasp movements, several motion parameters were calculated from the motion tracking data. Reaches that were initiated before the go-signal, or with both forward and backward movements were excluded from analysis (10 reaches, 0.24%, in Experiment 1 and 19 reaches, 0.48%, in Experiment 2). Missing samples (0.03% in Experiment 1 and 0.02% in Experiment 2) in the motion tracking data were linearly interpolated, and the data was subsequently filtered with a 4^th^ order low-pass Butterworth filter, with a cut-off frequency of 5 Hz. The data was differentiated with respect to time to obtain the velocity vector in three dimensions. The velocity value was calculated as the length of this vector, separately for each finger.

The different motion parameters are illustrated in Figure 1B. A reach-to-grasp movement could be described with a first, ballistic component and further secondary movements. Movement time was defined as the duration of the ballistic movement, between the time point the velocity of one of the fingers increased above a threshold of 5 cm/s (movement onset) and the time point the velocity of one of the fingers dropped again below 5 cm/s (movement offset). Peak velocity was calculated as the maximum velocity for the ballistic movement, i.e. between the time points that defined the ballistic phase, for thumb and index finger separately.

After the ballistic movement, one or more secondary movements could be made, especially in Experiment 2 where vision was not available (see Figure 1B). The secondary movement time was the time between the end of the ballistic movement and the start of the upward lifting movement. To define the start of the lifting movement, the point of a stable grasp was sought, which was the point at which the grip aperture (see below for calculation) differed less than 1 cm from the size of the object and stayed below this threshold for at least 0.5 s. The start of the lift was then defined as the time point in which a stable grasp was present and both fingers had a velocity above 5 cm/s in the vertical direction. Reach-to-grasp movements were checked by eye to ensure the correct point of lift start was found. For some movements, the point of stable grasp was manually adjusted (12 grasps in Experiment 1, 40 grasps in Experiment 2). Some lifts were very slow or absent and for those grasping movements, the vertical velocity criterion was decreased to 0 cm/s (286, including all 277 grasps of one participant that did not lift the object and 9 grasps of other participants in Experiment 1, 1 grasp in Experiment 2).

To evaluate the scaling of the reach-to-grasp movement to the size of the object and the change in object size, the grip aperture was calculated. The grip aperture was the distance between the markers on the thumb and index finger. Grip aperture values were corrected for the thickness of the fingers by subtracting the grip aperture value before movement onset, since participants always waited at the start position with thumb and index finger pinched together. The maximum grip aperture (MGA) was the largest grip aperture between movement onset and the start of the lifting movement.

Note that if multiple peaks in the grip aperture were seen due to corrections, the highest peak was used for the MGA. In addition, the MGA only reflects the grip aperture at one time point. Therefore, the MGA might not completely reflect the profile of the grip aperture in the reach-to-grasp movement. Therefore, another parameter was calculated that reflected the overall grip aperture across the reaching movement: the summed grip aperture. To adjust for differences in the duration of the reaching movement, the grip aperture was plotted against the relative reaching time between movement onset and start of the lifting movement, i.e. from 0-100% of the reaching time, including both movement time and correction time (see Figure 4). That is, the grip aperture was resampled to obtain 101 steps (0-100%). Next, these values were summed (summed grip aperture) to obtain a measure of the grip aperture profile over the course of the reach. The summed grip aperture can be illustrated as the area under the curve, as visualised in Figure 1B. Note that in this figure the x-axis is not shown as a relative measure since only a single reaching movement is shown.

**Figure 3.**
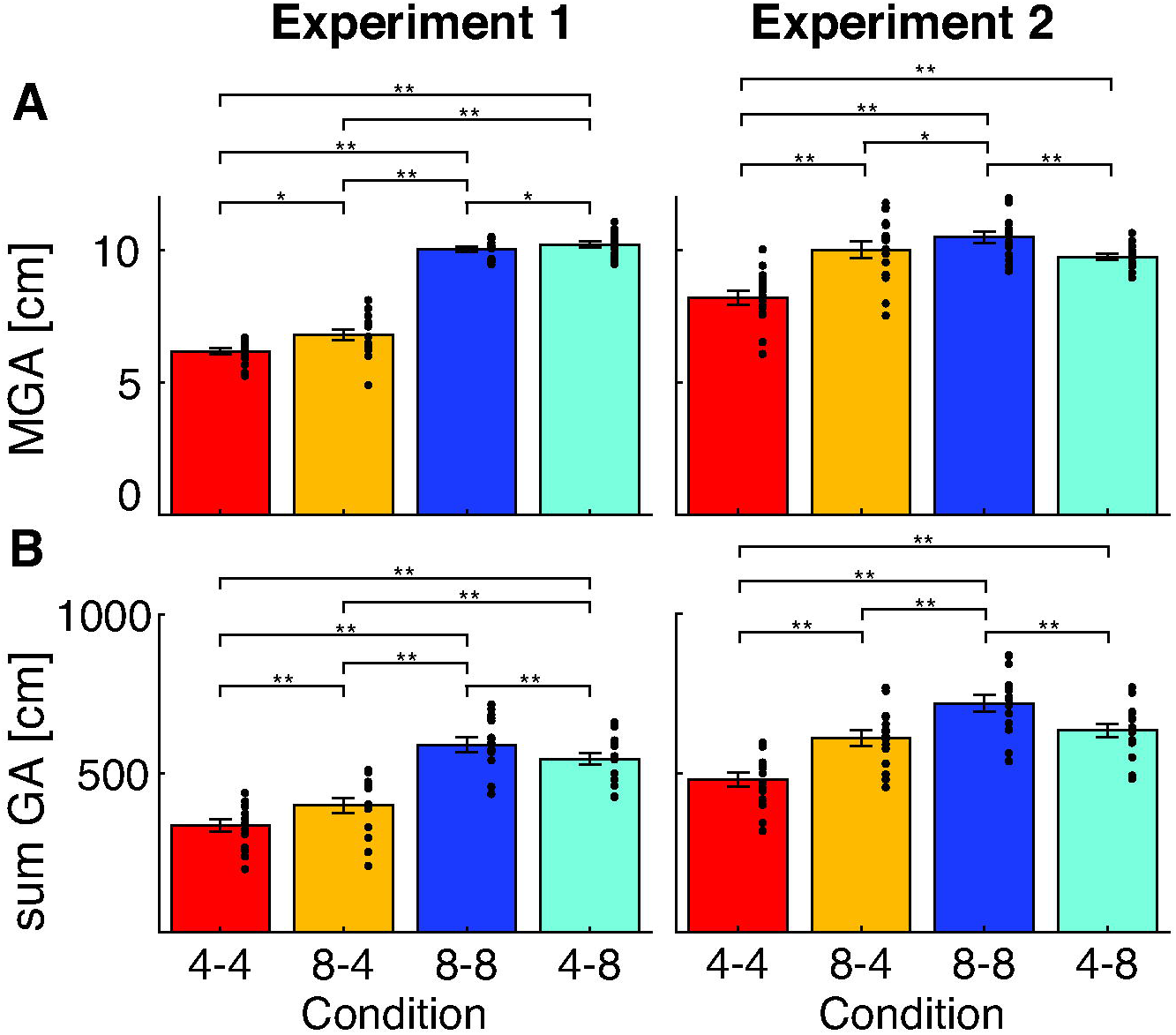
Grip aperture parameters for Experiment 1 (left) and Experiment 2 (right). **A:** Maximum grip aperture (MGA), and **B:** summed grip aperture (sum GA). Error bars represent standard errors, black dots indicate individual participant values. Significant differences are indicated with asterisks (*p<0.05, **p<0.001).

**Figure 4.**
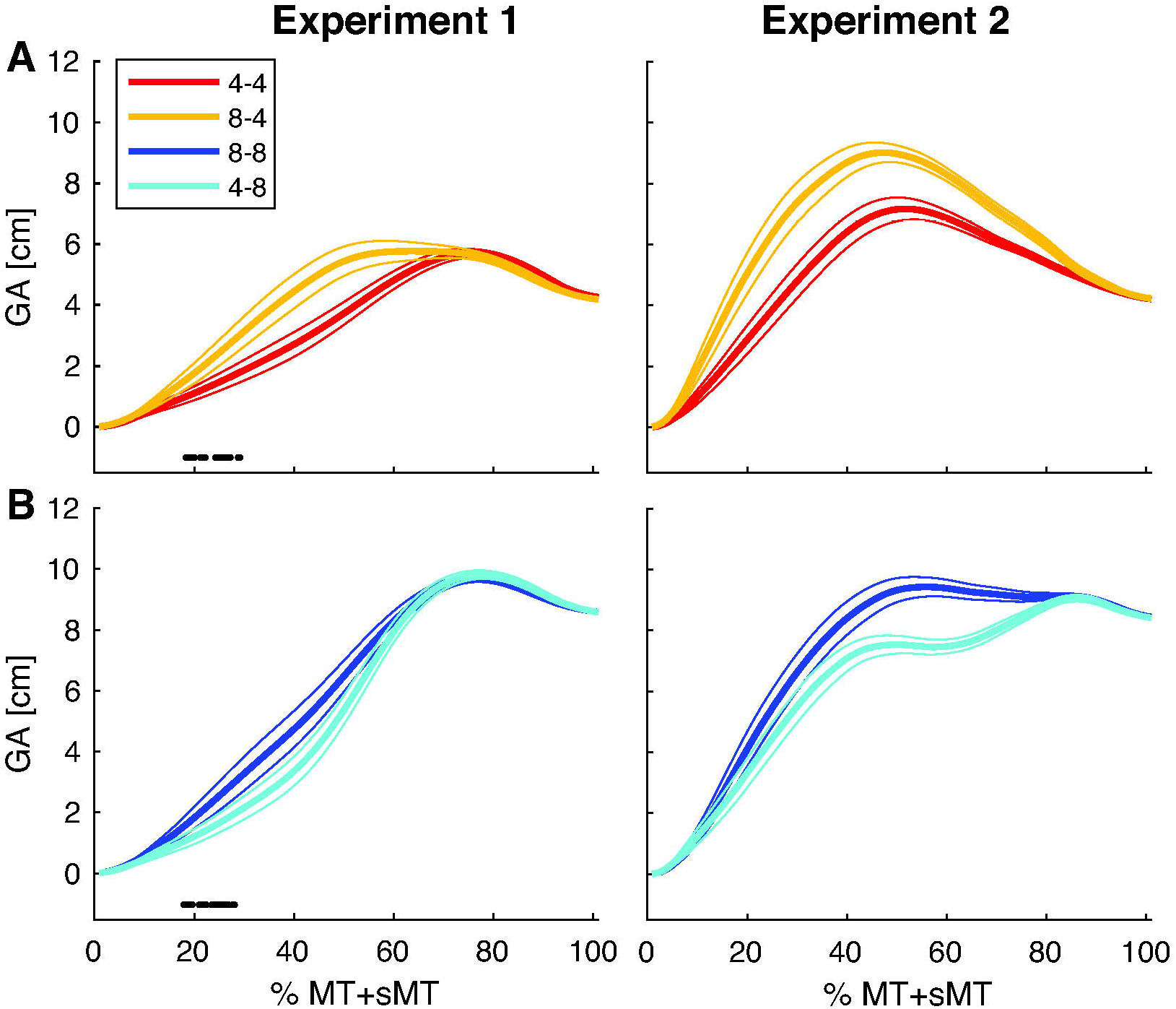
Grip aperture (GA) profiles. **A:** Grasping a small object with a small preview (red lines: no-correct small 4-4) or large preview (yellow: correct small 8-4). **B:** Grasping a large object with a large preview (blue: no-correct large 8-8) or small preview (cyan: correct large 4-8). Curves are plotted against the relative time from movement onset to the starting of the lifting movement, including both movement time (MT) and secondary movement time (sMT). Thick lines indicate average curves across participants, thin lines represent standard errors. Black dots represent individual time points the glasses opened in Experiment 1. These curves were used to calculate the summed grip aperture.

### 2.5 Statistics

Statistics were performed with SPSS (Version 27, IBM). One-sample t-tests were used to determine whether biases were different from zero. In addition, the biases and the weber fractions in the four conditions were compared in a 2 (size: large or small) × 2 (change: correct or no-correct) repeated measures analysis of variance (ANOVA).

For all movement parameters, only the reach-to-grasp movements towards the reference objects were used for statistical analysis. There were not enough reaches for each test object for statistical analysis. Furthermore, to compare performance between the different conditions, comparing reaches towards reference objects would be sufficient. All movement parameters (movement time, secondary movement time, peak velocity, MGA, summed grip aperture) were analysed with a 2 (size) × 2 (change) ANOVA as well.

For post-hoc tests, paired samples t-tests (two-tailed) with a Bonferroni correction were used. The alpha level was set to 0.05.

## RESULTS

In this study, participants performed reach-to-grasp movements for differently sized bars. They grasped two bars and responded which of the two was larger. They initiated their reaching movement without vision, but could plan their movement based on a preview of the object. Importantly, in some reaches the object increased or decreased in size after the preview, so participants had to correct their planned movement. In Experiment 1, vision was restored during the reaching movement, so visual information could be used to correct the movement. In Experiment 2, no visual information was available and corrections could only be based on haptic information. The aim of the experiments was to examine whether movement corrections would affect the perceived size of the grasped objects. In the results, values are represented as mean±standard error. All data used for analysis and figures are available at https://osf.io/ptukn/.

### 2.6 Grip aperture is adjusted when object size changes

First of all, it was investigated whether the experimental paradigm was successful in inducing corrections in the planned movements when the object changed in size between the preview and the grasp. If an object increases in size, the grip aperture needs to be increased, whereas a decrease in object size requires a reduction of the grip aperture.

As a first measure, the maximum grip aperture was calculated. This MGA is usually scaled towards the expected size of the object [1]. Results for the MGA are shown in Figure 3A. As expected, in Experiment 1, the MGA was larger when reaching for large objects compared to small ones (main effect size, F(1,14)=883.8, p<0.001, *η*_p_^2^=0.98). Interestingly, it was also larger when objects changed in size (main effect change, F(1,14)=16.2, p=0.001, *η*_p_^2^=0.54) and there was a significant interaction between size and change (F(1,14)=4.7, p=0.048, *η*_p_^2^=0.25). Post-hoc tests indicated that all conditions differed from each other (all t>3.2, all p<0.042). Overall, these results suggest that the MGA was larger when reaching for large objects compared to small (no-correct small: 6.18±0.12 cm, no-correct large: 10.05±0.10 cm). In addition, a slightly larger MGA was observed when objects changed in size, both when grasping the small (6.81±0.21 cm) and large object (10.21±0.12 cm). However, the direction of the change did not affect the MGA: corrections to the grip aperture always resulted in an increase compared to no-correct conditions. This is surprising, since changes from small to large or vice versa require corrections in opposite directions. However, the MGA only provides the grip aperture at a specific time point and is always the maximum value. That is, if the grip aperture changes from small to large, or from large to small, in both cases the MGA would be the largest grip aperture.

To address this issue, also the complete grip aperture profile was evaluated, which is shown in Figure 4. To adjust for the reaching time (including both the movement time and the secondary movement time), the grip aperture was resampled and plotted against the relative reaching time (0-100%). In Figure 4 can be clearly seen that the grip aperture is different depending on the preview of the object. When grasping a small object (Figure 4A), the grip aperture is initially larger with a preview of a large object (yellow line) compared to a preview of a small object (red line). The initially planned large aperture then has to be reduced to a small aperture in order to grasp the object. Similarly, when grasping a large object, the grip aperture is initially smaller when there was a preview of a small object (cyan line) compared to a preview of a large object (blue line). The initial small aperture is then quickly increased to a large aperture to grasp the large object.

To quantify these grip aperture profiles and statistically compare the different conditions, the summed grip aperture was calculated from these profiles (Figure 3B). The ANOVA on the summed grip aperture in Experiment 1 showed significant effects of size (F(1,14)=772.6, p<0.001, *η*_p_^2^=0.98), change (F(1,14)=7.8, p=0.014, *η*_p_^2^=0.36) and an interaction (F(1,14)=28.3, p<0.001, *η*_p_^2^=0.67). Post-hoc comparisons indicated that all conditions differed from each other (all t>5.1, all p<0.001). The summed grip aperture was larger for large objects compared to small objects. Furthermore, when grasping a small object, the correct small condition (399.1±18.2 cm), with a preview of a large object, had a larger summed grip aperture than the no-correct small condition (336.6±18.2 cm), with a preview of a small object. Similarly, when grasping a large object, a large preview resulted in a larger summed grip aperture: the summed grip aperture was larger in the no-correct large (589.7±23.9 cm) than the correct large condition (545.9±19.0 cm). These results show that the preview affected the grip aperture profile, in the direction of the previewed size. When the preview was small and the object changed to large, the grip aperture was initially smaller and then increased, as seen in Figure 4, hence the summed grip aperture was smaller than the no-correct large condition. Similarly, when the preview was large in the correct small condition, the grip aperture was initially larger and then adjusted, resulting in a larger summed grip aperture than the no-correct small condition. This shows that the experimental manipulation was successful in adjusting the intended grasp corrections in Experiment 1 in opposite directions when changing from large to small or from small to large.

In Experiment 2, the same measures of MGA and summed grip aperture were calculated, which are also shown in Figure 3 and 4. The MGA also depended on object size and change (effect size: F(1,14)=34.0, p<0.001, *η*_p_^2^=0.71, effect change: F(1,14)=63.6, p<0.001, *η*_p_^2^=0.82) and these factors also interacted (F(1,14)=130.6, p<0.001, *η*_p_^2^=0.90). Post-hoc tests indicated that larger objects had a larger MGA when objects did not change (no-correct large, 10.47±0.21, vs. no-correct small, 8.20±0.27, t(14)=-15.6, p<0.001) and when comparing grasped objects with similar previews (no-correct large vs. correct small, 10.00±0.32, and correct large, 9.74±0.11, vs. no-correct small, both t>3.3, p<0.033). Beside this effect of object size, an object change affected the MGA as well. When reaches with a change were compared to reaches without change, conditions with a preview of a large object resulted in a larger MGA, for both grasped object sizes (correct small vs. no-correct small and correct large vs. no-correct large, both t>5.6, p<0.001). In other words, when a small object changed to a large one, MGA increased and when a large object changed to a small one, MGA decreased. Finally, the two correct conditions did not differ from each other in MGA (correct small vs. correct large).

The summed grip aperture showed a similar effect. There was a main effect of size (F(1,14)=490.8, p<0.001, *η*_p_^2^=0.97) and change (F(1,14)=27.1, p<0.001, *η*_p_^2^=0.66), but these should be interpreted in light of the interaction (F(1,14)=180.4, p<0.001, *η*_p_^2^=0.93). Reaches for small objects had a smaller summed grip aperture than those for large objects when comparing reaches without a change (no-correct large, 718.9±26.2, vs. no-correct small, 480.6±22.2), a small preview (correct large, 634.1±21.1, vs. no-correct small) and a large preview (no-correct large vs. correct small, 610.1±24.0) (all t>14.9, p<0.001). In addition, a change in object size influenced the summed grip aperture, where a large preview resulted in a larger summed grip aperture, both when grasping small (correct small, vs. no-correct small) and large objects (no-correct large vs. correct large) (both t>9.2, p<0.001). In other words, when the object increased in size, the summed grip aperture was smaller than without such an increase, and when the object decreased in size, the summed aperture was larger than without a decrease. Similar to the MGA, the summed grip aperture did not differ between both correct conditions (correct small vs. correct large).

To sum up, in both experiments, the grip aperture was affected by the size of the grasped object and by a change in object size. In Experiment 1, the size of the grasped object increased the MGA and any change in object size increased MGA. While the expected direction in object change was not visible in the MGA, it was seen in the summed grip aperture, where increases/decreases in object size resulted in smaller/larger summed grip apertures, reflecting the preview size. In Experiment 2, the effect of both the grasped object size and a change in object size from preview to grasp altered the grip aperture, which was similarly visible in the MGA and summed grip aperture. Here, the effects of object size and preview size seemed to cancel out each other, resulting in no differences between the two correct conditions. All in all, it seems that the experimental paradigm was successful in altering the grip apertures and inducing corrections towards smaller or larger objects when they changed in size. Next, it was evaluated whether these corrections altered the perceived size of the grasped objects.

### 2.7 Perceptual biases and discrimination performance

Participants sequentially grasped two objects and compared these in size. In the no-correct conditions, they compared two objects (a test and a reference) that both did not change in size. In the correct conditions, they compared a test object that did not change in size to a reference object that changed in size between a preparation and execution phase. Participants answered whether the test or reference object was larger. From these answers, the perceptual bias with respect to the reference object was calculated. The perceptual biases are shown in Figure 5. In panel A and B, the average psychometric curves are shown and biases are indicated with dashed lines. If the dashed line deviates from the solid black line, a perceptual bias is found. Individual biases are shown in panel C and E, where they are compared between correct and no-correct conditions.

**Figure 5.**
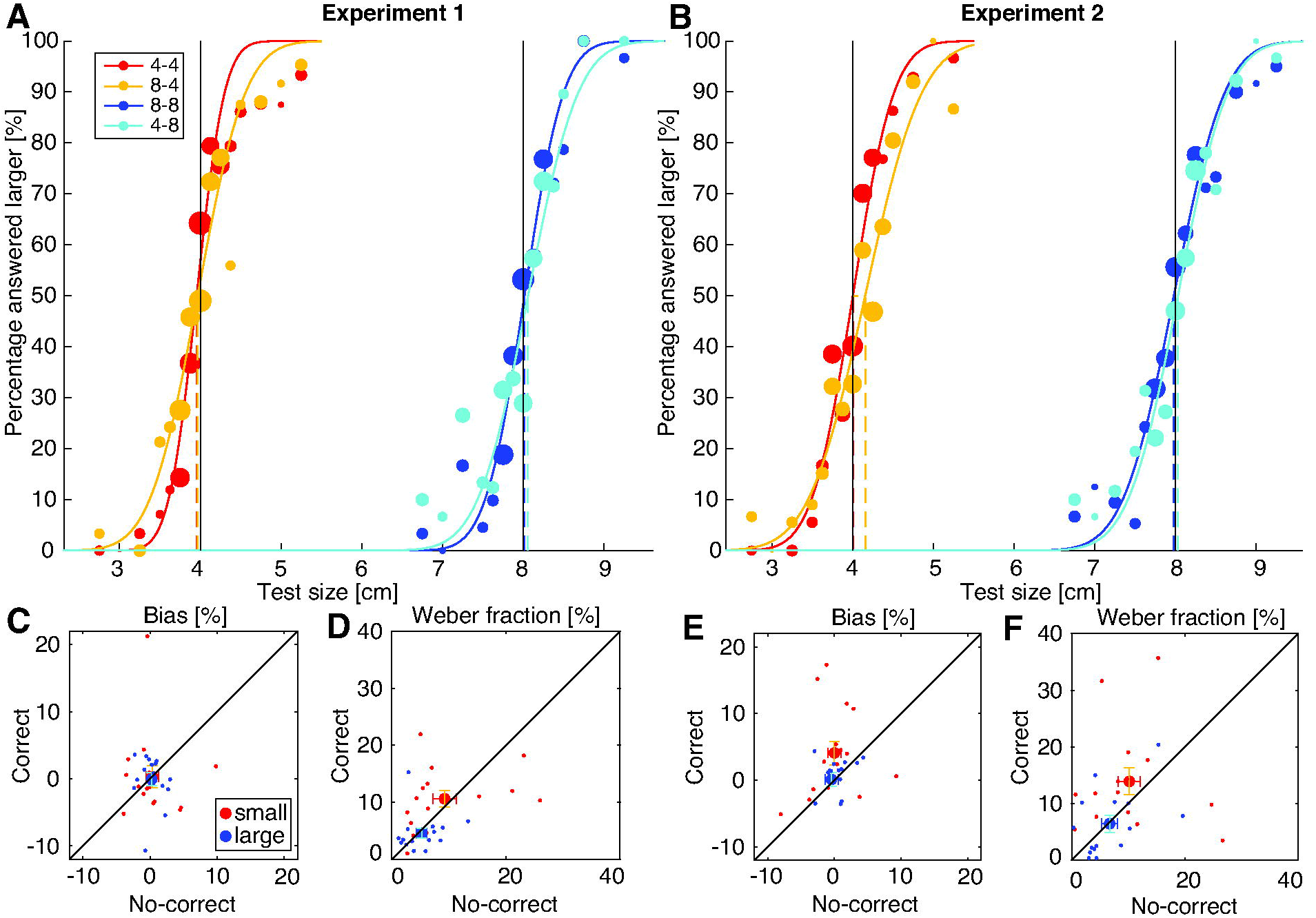
Perceptual biases and weber fractions. **A-B**: Average psychometrical curves in Experiment 1 (A) and 2 (B). Solid lines indicate reference values and dashed lines indicate biases. Dot size represents the number of trials. Colours indicate different conditions: no-correct small (4-4), correct small (8-4), no-correct large (8-8), correct large (4-8). **C-F:** Biases (C,E) and weber fractions (D,F) as obtained from the fitted curves for both experiments. Values from the correct conditions are plotted against those from the no-correct conditions, for a small and large reference separately. Error bars represent standard error, dots indicate individual participant values. The diagonal black line represents the points where the correct condition is equal to the no-correct condition. It can be seen that most values are scattered around the diagonal, indicating no differences between correct and no-correct conditions, except for the bias for small objects in Experiment 2. Weber fractions are on average larger for small compared to large objects in both experiments.

In Experiment 1, participants were able to view the object for the end part of the reach-to-grasp movement. Biases in all conditions were not significantly different from zero (all p>0.60). No differences were found between the different conditions either (no-correct small: 0.31±0.93%, correct small: 0.31±1.67%, no-correct large: 0.23±0.45%, correct large: −0.08±0.98%).

In Experiment 2, there was a main effect of change (F(1,14)=5.0, p=0.041, *η*_p_=0.27) that indicated that the bias was larger in conditions were the object changed in size. When testing the individual conditions, only the bias in the correct small condition was significantly different from zero (4.09±1.76%, p=0.035). The other biases did not differ from zero (no-correct small: 0.02±1.02%, no-correct large: −0.42±0.99%, correct large: 0.11±1.01%). This positive bias indicates that the grasped object is perceived to be smaller when it changes from a large preview to a small object.

The weber fractions of each condition can be found in Figure 5D and F. In Experiment 1, it can be seen that the weber fraction appears larger for the stimuli in the small set (no-correct small: 9.24±2.06%, correct small: 10.62±1.42%) than in the large set (no-correct large: 5.22±0.88%, correct large: 4.59±0.86%). This was confirmed by a 2 (size) × 2 (change) ANOVA, which indicated a significant effect of size (F(1,14)=12.4, p=0.003, *η*_p_^2^=0.47). This suggests that discrimination between the objects is easier for the large object set, regardless of a change in size during grasping. Similarly, in Experiment 2, the weber fraction was smaller for large objects (no-correct large: 6.69±1.37, correct large: 6.43±1.51) compared to small objects (no-correct small: 10.19±1.96, correct small: 13.92±2.36; effect size, F(1,14)=16.2, p=0.001, *η*_p_^2^=0.54). No effects of change were found in neither experiment.

The biases were smaller than the weber fractions in both experiments. This was confirmed by paired samples t-tests between the biases and the weber fractions (all conditions p<0.015). This suggests that the minimal difference in size that can be perceived is larger than any shift in size perception induced by changing the objects. In the study of Cesanek and Domini (21), an effect of grasp adaptation on manual estimation of ~4 mm was found. With an average size of 44 mm of the objects in their experiment, this would suggest a bias of 10%. The biases in the present experiments were all significantly smaller than this value (one-sample t-tests, all p<0.013).

### 2.8 Correlations between perceptual bias and grip aperture

Since there was a large variability in biases and grip aperture for different participants, it was examined whether effects on these parameters could be related. That is, the hypothesis would suggest that a larger effect on the grip aperture would result in a larger perceptual bias. The correlation between the perceptual effect (difference in bias, e.g. correct small – no-correct small) and the grasping effect (difference in summed grip aperture, e.g. correct small – no-correct small) was calculated for both object sizes. In Experiment 1, correlation coefficients of – 0.07 (p=0.79) and −0.18 (p=0.51) were found for the small and large reference, respectively. In Experiment 2, correlation coefficients for the small and large reference were 0.26 (p=0.36) and −0.06 (p=0.83), respectively. Even if both experiments were combined to obtain more data points, correlations were still not significant: 0.22 (p=0.24) and −0.17 (p=0.36) for a small and large reference. Therefore, none of these correlations were significant.

### 2.9 Reaching parameters: Movement Time and peak velocity

Finally, to see whether the experimental conditions differed in other aspects of the reaching movement besides the grip aperture, standard measures such as the movement time of the ballistic movement and secondary movements, and the peak velocity of both fingers were calculated.

Movement times of the ballistic movement in Experiment 1 were 1.223±0.042 s (no-correct small condition), 1.219±0.039 s (correct small), 1.221±0.039 s (no-correct large) and 1.254±0.041 s (correct large). The movement time in Experiment 1 was shorter for small objects than for large objects (main effect size, F(1,14)=9.4, p=0.008, *η*_p_^2^=0.40). An interaction of size and change (F(1,14)=17.3, p=0.001, *η*_p_^2^=0.55) indicated that this difference between object size was only seen when objects changed in size (i.e. correct small vs. correct large, t(14)=-5.2, p<0.001). In addition, when grasping a large object, movements were slower when the object changed, but not when it stayed the same as the preview (i.e. correct large vs. no-correct large, t(14)=-3.8, p=0.012).

In Experiment 2, movement times were 1.172±0.043 s (no-correct small), 1.246±0.047 s (correct small), 1.162±0.043 s (no-correct large) and 1.134±0.047 s (correct large). Similar to the 1^st^ experiment, movement time also depended on object size (F(1,14)=11.9, p=0.004, *η*_p_^2^=0.46), but here the grasping of large objects was faster than small objects. There was also a main effect of change (F(1,14)=5.4, p=0.036, *η*_p_^2^=0.28) and an interaction of size and change (F(1,14)=22.8, p<0.001, *η*_p_^2^=0.62). Post-hoc tests indicated that the change from a large to a small object (correct small) had a longer movement time than all other conditions.

Results for the secondary movement time in Experiment 1, only showed a main effect of size (F(1,14)=26.6, p<0.001, *η*_p_^2^=0.66). Here, the time was slightly longer for large objects (0.28±0.04 s) compared to small objects (0.24±0.04 s).

In Experiment 2, the secondary movement time was affected by a change in object size (F(1,14)=115.5, p<0.001, *η*_p_^2^=0.89), where a longer time was seen when the preview changed. There was also an interaction of change and size (F(1,14)=5.4, p=0.036, *η*_p_^2^=0.28). Post-hoc tests showed that the conditions in which a change was present (correct small: 1.07±0.08 s, correct large: 0.98±0.05 s), times were longer than all other conditions (no-correct small: 0.76±0.07 s, no-correct large: 0.78±0.06 s; all t>4.7, all p<0.002), but not different from each other.

Finally, the peak velocity was calculated separately for the thumb and index finger. No differences between the conditions were found in peak velocity of the thumb in both experiments (grand mean 83.1±3.92 cm/s and 86.3±4.35 cm/s for Experiment 1 and 2, respectively).

For the velocity of the index finger, only an interaction between size and change was found (F(1,14)=12.9, p=0.003, *η*_p_^2^=0.48) in Experiment 1. For each condition, peak velocity was 85.88±3.98 cm/s (no-correct small condition), 87.30±4.05 cm/s (correct small), 88.11±4.02 cm/s (no-correct large), and 86.26±4.12 cm/s (correct large). Post-hoc tests indicated that the velocity was larger when grasping a large object that did not change in size compared to a large object that did change in size (no-correct large vs. correct large, t(14)=-3.3, p=0.030). In Experiment 2, also an interaction between size and change was found (F(1,14)=8.6, p=0.011, *η*_p_^2^=0.38) for peak velocity of the index finger. Peak velocities were 91.47±4.51 cm/s (no-correct small condition), 91.79±4.44 cm/s (correct small), 93.14±4.56 cm/s (no-correct large) and 90.54±4.66 cm/s (correct large). Here, it was also found that the velocity was larger when reaching for large objects that did not change in size (no-correct large) compared to a change in size (correct large) (t(14)=3.1, p=0.046).

All in all, there seemed to be only small differences in movement time and peak velocity with a change in object size. The movement time was only increased for large objects in Experiment 1 and only for small objects in Experiment 2 and the differences between the conditions in movement time appeared to be small. Peak velocities were not altered for the thumb, but lower for the index finger when objects changed from a small to large size. The secondary movement time was small and not altered by a change in object size in Experiment 1. On the other hand, in Experiment 2, longer secondary movement times were seen when the object changed in size. Since participants could not see the object in Experiment 2, they could only detect a change in object size after contacting the object and then needed time to adjust their grip before commencing the lift.

## 3 DISCUSSION

The present study investigated whether corrections to planned grasping movements (i.e. grip aperture) affected the size perception of grasped objects. The grip aperture was altered by changing the planned movement with a preview object that could sometimes change to an object of a smaller or larger size after the reach-to-grasp movement was initiated. In case of a change, the planned movement, and the grip aperture, had to be corrected during the reach. While the experimental paradigm was successful in altering grip apertures, this did not result in a biased size perception. Therefore, the planning and execution of grasping movements does not seem to affect the perception of object size.

In the experiment, participants corrected their grip aperture to the new object if this was different from the preview size. The grip aperture was increased if the object became larger and was decreased when the object became smaller. In case of a change in object size, the movement was somewhat slower, as seen in longer movement times and lower peak velocities, but otherwise the reach-to-grasp movement was quite similar to conditions without a change. These findings agree with previous studies that found smooth adaptations of grip aperture to a change in object size during reach [26, 27].

The experimental task was memory-demanding, requiring participants to remember the size of the first grasped object, and not the previewed size of the first trial, and compare this with the grasped object of the second trial, while ignoring the size of the preview of the second trial. Despite this complicated paradigm, participants seemed to be quite good at comparing the grasped objects, without being confused by the previewed objects. However, they could use the preview to plan their grasping movements, which would then need to be altered if the object changed in size. Despite appropriate changes in grasping movements, no significant perceptual biases were found in Experiment 1. In Experiment 2, only a significant perceptual bias was found when the object changed from large to small, but not from small to large. In addition, this bias was much smaller (4.1%) than the weber fraction (13.9%). The weber fraction indicates the minimal difference in size that can be reliably perceived, i.e. the discrimination threshold. Since the bias was smaller than this threshold, it would not have been perceivable by the participants. Previous research found a smaller discrimination threshold of 4% for size perception [2], but this was with a combination of visual and haptic information. If only haptic information was available, as in Experiment 2, a threshold of about 9% was found [2], which is still larger than the bias found in Experiment 2. All in all, although the change in object size altered the grip aperture, this did not lead to a meaningful difference in size perception.

Recent studies suggested that altered grip apertures could affect the manual estimation of size [21] or the estimated graspability of the object [20]. The former study used an adaptation paradigm in virtual reality to alter the grip aperture. It is possible that only such a long-term protocol would be effective for altering size perception, while this would not be found on a trial-by-trial basis when grip aperture is only adjusted for a single trial. In addition, while visual size-perception was unaltered in their study, it is possible that the visual-haptic integration was altered by the adaptation. A different relation between vision and proprioception due to the adaptation might have possibly led to altered manual size estimations. The other study [20] only found effects of grip aperture alterations on graspability of objects, not on size perception. Since only size perception was measured in the present study, it seems that a judgement that is more related to movement kinematics, i.e. graspability, would be more easily influenced by an action.

It was hypothesized that specifically corrections to an action plan would alter perception. Such an effect was found in object lifting studies [22, 23], where the erroneously planned forces or torques were associated with differences in perceived weight. When the weight or weight distribution was different than expected, forces or torques had to be corrected and the estimated weight of the object was altered as well. Furthermore, the effect on the action parameters and the perceptual estimates correlated (although it must be noted that a recent study did not find a correlation between the effects on force scaling and weight perception [28]). In the present study, corrections were seen on the grip aperture in a reach-to-grasp movement, but no differences in size perception were seen. Therefore, it seems that interactions between action and perception might be different for different tasks. Different control mechanisms might exist for object grasping and object lifting movements. Possibly, the link between force scaling and weight perception is stronger than the link between grasp scaling and size perception. One theory of grasp control suggests that the distance between the fingertips and the position of the hand is controlled to successfully grasp objects of different sizes [1, 29, 30]. Since the opening of the fingers has to be shaped to the size of the object, this could suggest a relation to the perception of object size. However, the present experiment and others suggest that these two processes are not necessarily related. For instance, previous studies found that illusions can have different effects on size perception and grasp shaping ([4, 5] but also see [6]). The results could also be interpreted in view of another theory of grasp control, where the digits are pointing to the grasp positions on the object [31, 32]. According to this view, the grasping plan is not based on the size of the object, but on the positions the fingers will grasp the object. In this way, the grasping movement might not be related to the perception of size, because different sources of information are used to control these processes, i.e. size and position.

A second aim of this study was to evaluate the role of visual information in the relation between grasp corrections and size perception. In the first experiment, visual and haptic information was available to correct grasping movement, whereas in the second experiment only haptic information was available. In Experiment 1, corrections could be initiated earlier than in Experiment 2, hence smaller corrections were needed in the first experiment. In the first experiment, participants were able to see the object they grasped. It was possible that participants only relied on the object size that they had to grasp and ignored the preview. In fact, while most participants initially planned their grip aperture based on the preview object and corrected this grip aperture during the reach if the object changed, there were a few participants that did not show this behaviour. These participants kept their fingers pressed together during reach until the glasses opened and only then scaled their grip aperture to the size of the object to be grasped. Specifically, in the first experiment, the grasping of the object was not necessary to decide whether the object differed in size from the previous trial, because vision was available at the end. Therefore, the second experiment was performed in which vision was blocked throughout the reach-to-grasp movement. In this way, the object needed to be grasped to perceive the object size and to be able to compare this size to the previously grasped object. Although this setup appeared to have a larger effect on the grip aperture corrections, the effects on the size perception were small. Hence, the perception of object size was largely unaffected by the availability of vision.

Since the perceptual estimates did not seem to be altered much by the availability of visual information, it is likely that participants mostly relied on haptic information for their size judgements. In this study, there was no condition with only visual and no haptic information. This might have been accomplished by letting participants pantomime the grasp. However, there is evidence indicating that pantomiming grasping movements differ from actual grasping movements [33], especially when no haptic feedback is provided [34], which would cast doubt on the results, which would perhaps only be valid when pantomiming grasps. Future studies could shed further light on the potential roles of haptic perception and pantomimed grasps in size perception.

A final note about the current experiment can be made regarding the discrimination of object size. It was found that the weber fractions of large objects were smaller than those for small objects. This finding was unexpected, because often the discrimination threshold of perceived properties depends on the intensity of the object. In other words, for larger objects, a larger difference is needed to be able to be perceived. If the threshold is expressed in a relative measure, e.g. the weber fraction, this measure is usually constant for perceived properties. Interestingly, this does not hold for grasping parameters, where variability does not seem to increase with object size [35]. In the current experiment, the absolute perceptual thresholds seemed to be constant in the small and large set (about 4-5 mm). One study that investigated a large range of object sizes found that variability of perceptual estimations increased with object sizes, but up to a certain point where it remained constant [36]. This was possibly related to biomechanical constraints when grasping large objects. Since the present study used relatively large object sizes, it is possible that the variability was close to a ceiling point and did not further increase with size. Importantly, although the weber fraction was different for large and small objects, it was not altered by the change in object size, indicating that the grasp aperture correction did not influence the perceptual discrimination threshold.

In conclusion, it was investigated whether corrections to grip apertures during grasping would influence the perceived size of the grasped object. No such an effect was seen when objects were visible during grasping and only a small effect was seen when objects were not visible. However, this effect was smaller than the size difference that can be reliably perceived and does not seem to be a meaningful effect on size perception. Therefore, it can be concluded that the planning and execution of grasping movements do not reliably affect size perception of objects.

## ACKNOWLEDGEMENTS

The author would like to thank Jean-Jacques Orban de Xivry for helpful comments on an earlier version of this paper, and Arsalis (Belgium) for providing the stimulus bars.

## REFERENCES

1. Jeannerod M, Arbib MA, Rizzolatti G, Sakata H. Grasping objects: the cortical mechanisms of visuomotor transformation. Trends Neurosci. 1995;18(7):314–20. doi: 10.1016/0166-2236(95)93921-J.

2. Ernst MO, Banks MS. Humans integrate visual and haptic information in a statistically optimal fashion. Nature. 2002;415(6870):429–33. doi: 10.1038/415429a.

3. Goodale MA, Milner AD. Seperate visual pathways for perception and action. Trends Neurosci. 1992;15:20–5.

4. Aglioti S, DeSouza JFX, Goodale MA. Size-contrast illusions deceive the eye but not the hand. Curr Biol. 1995;5(6):679–85. doi: 10.1016/s0960-9822(95)00133-3.

5. Haffenden AM, Goodale MA. The effect of pictorial illusion on prehension and perception. J Cogn Neurosci. 1998;10(1):122–36. doi: 10.1162/089892998563824.

6. Smeets JBJ, Brenner E. 10 years of illusions. J Exp Psychol Hum Percept Perform. 2006;32(6):1501–4. doi: 10.1037/0096-1523.32.6.1501.

7. Franz VH, Gegenfurtner KR. Grasping visual illusions: Consistent data and no dissociation. Cogn Neuropsychol. 2008;25(7):920–50. doi: 10.1080/02643290701862449.

8. Kopiske KK, Bruno N, Hesse C, Schenk T, Franz VH. The functional subdivision of the visual brain: Is there a real illusion effect on action? A multi-lab replication study. Cortex. 2016;79:130–52. doi: 10.1016/j.cortex.2016.03.020.

9. Pavani F, Boscagli I, Benvenuti F, Rabuffetti M, Farne A. Are perception and action affected differently by the Titchener circles illusion? Exp Brain Res. 1999;127(1):95–101. doi: 10.1007/s002210050777.

10. Franz VH, Hesse C, Kollath S. Visual illusions, delayed grasping, and memory: no shift from dorsal to ventral control. Neuropsychologia. 2009;47(6):1518–31. doi: 10.1016/j.neuropsychologia.2008.08.029.

11. Glover S. Visual illusions affect planning but not control. Trends in Cognitive Sciences. 2002;6(7):288–92. doi: 10.1016/s1364-6613(02)01920-4.

12. Vishton PM, Rea JG, Cutting JE, Nunez LN. Comparing effects of the horizontal-vertical illusion on grip scaling and judgment: relative versus absolute, not perception versus action. J Exp Psychol Hum Percept Perform. 1999;25(6):1659–72. doi: 10.1037//0096-1523.25.6.1659.

13. Schenk T, McIntosh RD. Do we have independent visual streams for perception and action? Cogn Neurosci. 2010;1(1):52–62. doi: 10.1080/17588920903388950.

14. Cloutman LL. Interaction between dorsal and ventral processing streams: where, when and how? Brain Lang. 2013;127(2):251–63. doi: 10.1016/j.bandl.2012.08.003.

15. van Polanen V, Davare M. Interactions between dorsal and ventral streams for controlling skilled grasp. Neuropsychologia. 2015;79(Pt B):186–91. doi: 10.1016/j.neuropsychologia.2015.07.010.

16. Gutteling TP, Kenemans JL, Neggers SF. Grasping preparation enhances orientation change detection. PLoS One. 2011;6(3):e17675. doi: 10.1371/journal.pone.0017675.

17. Wykowska A, Maldonado A, Beetz M, Schubö A. How Humans Optimize Their Interaction with the Environment: The Impact of Action Context on Human Perception. Int J Soc Robot. 2010;3(3):223–31. doi: 10.1007/s12369-010-0078-3.

18. Linkenauger SA, Witt JK, Proffitt DR. Taking a hands-on approach: apparent grasping ability scales the perception of object size. J Exp Psychol Hum Percept Perform. 2011;37(5):1432–41. doi: 10.1037/a0024248.

19. Collier ES, Lawson R. Does grasping capacity influence object size estimates? It depends on the context. Atten Percept Psychophys. 2017;79(7):2117–31. doi: 10.3758/s13414-017-1344-3.

20. Geers L, Pesenti M, Andres M. Visual illusions modify object size estimates for prospective action judgements. Neuropsychologia. 2018;117:211–21. doi: 10.1016/j.neuropsychologia.2018.06.003.

21. Cesanek E, Domini F. Transfer of adaptation reveals shared mechanism in grasping and manual estimation. Neuropsychologia. 2018;117(March):271–7. doi: 10.1016/j.neuropsychologia.2018.06.014.

22. van Polanen V, Davare M. Sensorimotor Memory Biases Weight Perception During Object Lifting. Front Hum Neurosci. 2015;9:700. doi: 10.3389/fnhum.2015.00700.

23. Schneider TR, Buckingham G, Hermsdorfer J. Torque-planning errors affect the perception of object properties and sensorimotor memories during object manipulation in uncertain grasp situations. J Neurophysiol. 2019;121(4):1289–99. doi: 10.1152/jn.00710.2018.

24. Paulignan Y, Jeannerod M, Mackenzie C, Marteniuk R. Selective perturbation of visual input during prehension movements. 2. The effects of changing object size. Exp Brain Res. 1991;87(2):407–20.

25. Camponogara I, Volcic R. Grasping adjustments to haptic, visual, and visuo-haptic object perturbations are contingent on the sensory modality. J Neurophysiol. 2019;122(6):2614–20. doi: 10.1152/jn.00452.2019.

26. Hesse C, Franz VH. Corrective processes in grasping after perturbations of object size. J Mot Behav. 2009;41(3):253–73. doi: 10.3200/JMBR.41.3.253-273.

27. van de Kamp C, Bongers RM, Zaal FT. Effects of changing object size during prehension. J Mot Behav. 2009;41(5):427–35. doi: 10.3200/35-08-033.

28. van Polanen V, Rens G, Davare M. The role of the anterior intraparietal sulcus and the lateral occipital cortex in fingertip force scaling and weight perception during object lifting. J Neurophysiol. 2020;124(2):557–73. doi: 10.1152/jn.00771.2019.

29. Jeannerod M. The neural and behavioural organization of goal-directed movements: Oxford: Clarendon; 1988.

30. Jeannerod M. Intersegmental coordination during reaching at natural visual objects. In: Long J, Baddely A, editors. Attention and Performance IX. Hillsdale, NJ: Erlbaum; 1981. p. 153–68.

31. Smeets JBJ, Brenner E. A new view on grasping. Motor control. 1999;3(3):237–71.

32. Smeets JBJ, van der Kooij K, Brenner E. A review of grasping as the movements of digits in space. J Neurophysiol. 2019;122(4):1578–97. doi: 10.1152/jn.00123.2019.

33. Goodale MA, Jakobson LS, Keillor JM. Differences in the visual control of pantomimed and natural grasping movements. Neuropsychologia. 1994;32(10):1159–78. doi: 10.1016/0028-3932(94)90100-7.

34. Davarpanah Jazi S, Yau M, Westwood DA, Heath M. Pantomime-grasping: the ‘return’ of haptic feedback supports the absolute specification of object size. Exp Brain Res. 2015;233(7):2029–40. doi: 10.1007/s00221-015-4274-0.

35. Ganel T, Chajut E, Algom D. Visual coding for action violates fundamental psychophysical principles. Curr Biol. 2008;18(14):R599–601. doi: 10.1016/j.cub.2008.04.052.

36. Bruno N, Uccelli S, Viviani E, de’Sperati C. Both vision-for-perception and vision-for-action follow Weber’s law at small object sizes, but violate it at larger sizes. Neuropsychologia. 2016;91:327–34. doi: 10.1016/j.neuropsychologia.2016.08.022.

